# AAV-mediated expression of proneural factors stimulates neurogenesis from adult Müller glia *in vivo*

**DOI:** 10.1101/2024.09.13.612930

**Authors:** Marina Pavlou, Marlene Probst, Elizaveta Filippova, Lew Kaplan, Aric R. Prieve, Fred Rieke, Thomas A. Reh

## Abstract

The lack of regeneration in the human central nervous system (CNS) has major health implications. To address this, we previously used transgenic mouse models to show that neurogenesis can be stimulated in the adult mammalian retina by driving regeneration programs that other species activate following injury. Expression of specific proneural factors in adult Müller glia causes them to re-enter the cell cycle and give rise to new neurons following retinal injury. To bring this strategy closer to clinical application, we now show that neurogenesis can also be stimulated when delivering these transcription factors to Müller glia using adeno-associated viral (AAV) vectors. AAV-mediated neurogenesis phenocopies the neurogenesis we observed from transgenic animals, with different proneural factor combinations giving rise to distinct neuronal subtypes *in vivo*. Vector-borne neurons are morphologically, transcriptomically and physiologically similar to bipolar and amacrine/ganglion-like neurons. These results represent a key step forward in developing a cellular reprogramming approach to regenerative medicine in the CNS.

## Introduction

According to the World Health Organization around 2.2 billion people worldwide suffer from vision impairment, caused largely by diseases such as age-related macular degeneration and glaucoma that are manifested by the death of cone photoreceptors and retinal ganglion cells (RGCs), respectively ^1^. This has sparked global efforts to develop therapies that can treat retinal degeneration, with some success in the small molecule, gene supplementation and gene editing fields for specific diseases ^2–6^. In most cases vision loss is caused by the irreversible death of key neuronal classes, specifically cone photoreceptors and RGCs. However, some organisms, like fish and amphibians, are capable of regenerating their retina ^7–9^. In fish, for example, the resident Müller glia (MG) upregulate key developmental factors in response to injury as they re-enter the cell cycle and dedifferentiate into multipotent progenitors that regenerate neurons and restore tissue function ^10–12^.

Recently, there has been significant progress towards inducing neurogenesis in the adult mammalian retina by mimicking natural regeneration pathways from other species. Using transgenic mouse models to drive the expression of developmental programs in adult MG can reprogram these cells into proliferating neurogenic progenitors and neurons. Moreover, different combinations of transcription factors (TFs) can stimulate MG to generate distinct neuronal lineages. For example, expression of achaete-scute family bHLH transcription factor 1 (Ascl1) alone converts MG into bipolar neurons, whereas combining Ascl1 with either atonal bHLH transcription factor 1 (Atoh1) or ISL LIM Homeobox 1 (Isl1) and POU domain class 4 transcription factor 2 (Pou4f2) converts MG into amacrine/retinal ganglion cells ^13–15^. Interestingly, the ability of mammalian MG to undergo proliferation and produce new neurons is not impacted by the timing or mode of injury, as transgenic expression of Ascl1 or Ascl1-Atoh1 will stimulate neurogenesis after excitotoxic injury or light-damage equally well, and the factors can be expressed before or after the injury with similar levels of neurogenesis ^16^.

Having identified combinations of factors that can push glial cells towards different neuronal lineages, the next challenge is to establish a way to translate this biology into a medical application for regenerative medicine. This requires the delivery of exogenous reprogramming factors in mammalian MG and driving their expression in a temporal manner. In this study, we investigate the feasibility of using viral vectors, specifically adeno-associated viral (AAV) vectors, to deliver the reprogramming factors to the MG and induce neurogenesis *in vivo*.

AAV vectors have become the gold standard for genetic perturbations in the eye, with several successful applications in the field of gene therapy against monogenic retinal disorders ^4–6,17,18^. This has led to several previous efforts to test AAV-mediated strategies to stimulate neurogenesis with proneural factors driven by cell specific promoters ^19^. Unfortunately, these early studies were confounded by the lack of glial-specificity of the promoters that were used, particularly when proneural TFs were expressed. This led to the false conclusion that AAV-mediated expression of neurogenic factors was inducing reprogramming of glia to neurons, when in fact the promoter specificity was altered by the expression of these TFs, leading to expression of the reporter in endogenous neurons, rather than the induction of neurogenesis ^19–24^. As such, we still lack convincing evidence that glia can proliferate and/or give rise to new neurons in response to AAV-borne expression of proneural factors.

To circumvent these artefacts, we used AAVs to deliver proneural factors in the adult mouse retina but used a transgenic mouse line to restrict the AAV payload expression to MG. Specifically, we used a transgenic mouse line (Rlbp1-CreERt2 x LSL-TdTomato) that enabled us to restrict Cre recombination only in MG in a tamoxifen-dependent manner, and thereby regulate the expression of the vector payload. In this way, the glial specificity is not conveyed by the AAV transgene cassette but rather by the transgenic line. This dual approach obviates the issues from the prior studies, but still allows us to test whether AAV delivered proneural factors can stimulate neurogenesis from MG in the adult mouse retina *in vivo*.

To maximize our targeting efficiency of MG in the adult mouse retina, we opted for the AAV.7m8 vector, which was shown to be potent when delivered through the vitreous ^25^. Using a combination of histology, EdU labelling of newborn cells, scRNA-seq and patch-clamp electrophysiology, we show for the first time that AAV-borne reprogramming factors can stimulate neurogenesis from MG after retinal injury. The efficiency of neurogenesis is affected by MG transduction, which is a function of both the vector titer and the incubation period of AAVs in the retina before inducing the expression cassette. Interestingly, the types of neurons generated by the MG are dictated by the transcription factors expressed and the genetic trajectory of glia-to-neuron conversion phenocopies our results from previously reported transgenic experiments. However, the physiological properties of neurons obtained after AAV-mediated neurogenesis were somewhat different than neurons obtained from transgenic mouse models of regeneration.

## Results

### Vector-borne Ascl1 expression stimulates MG proliferation and neurogenesis

In previous studies with transgenic mice, Ascl1 expression stimulated MG to re-enter the cell cycle and generate new bipolar neurons after inner or outer retinal injury and the subsequent administration of HDAC inhibitor Trichostatin-A (TSA) ^13,16^. To determine whether AAV-borne expression of Ascl1 in adult MG phenocopies this response, we injected adult mice intravitreally with AAV.7m8 vectors expressing an inducible FLip-EXcision (FLEX) cassette, where a ubiquitous promoter (CBh or Ef1α) is upstream the inverted Ascl1-TurboGFP cassette (Fig 1A). As a control, we used an equivalent vector expressing only the fluorescent reporter.

**Fig. 1.**
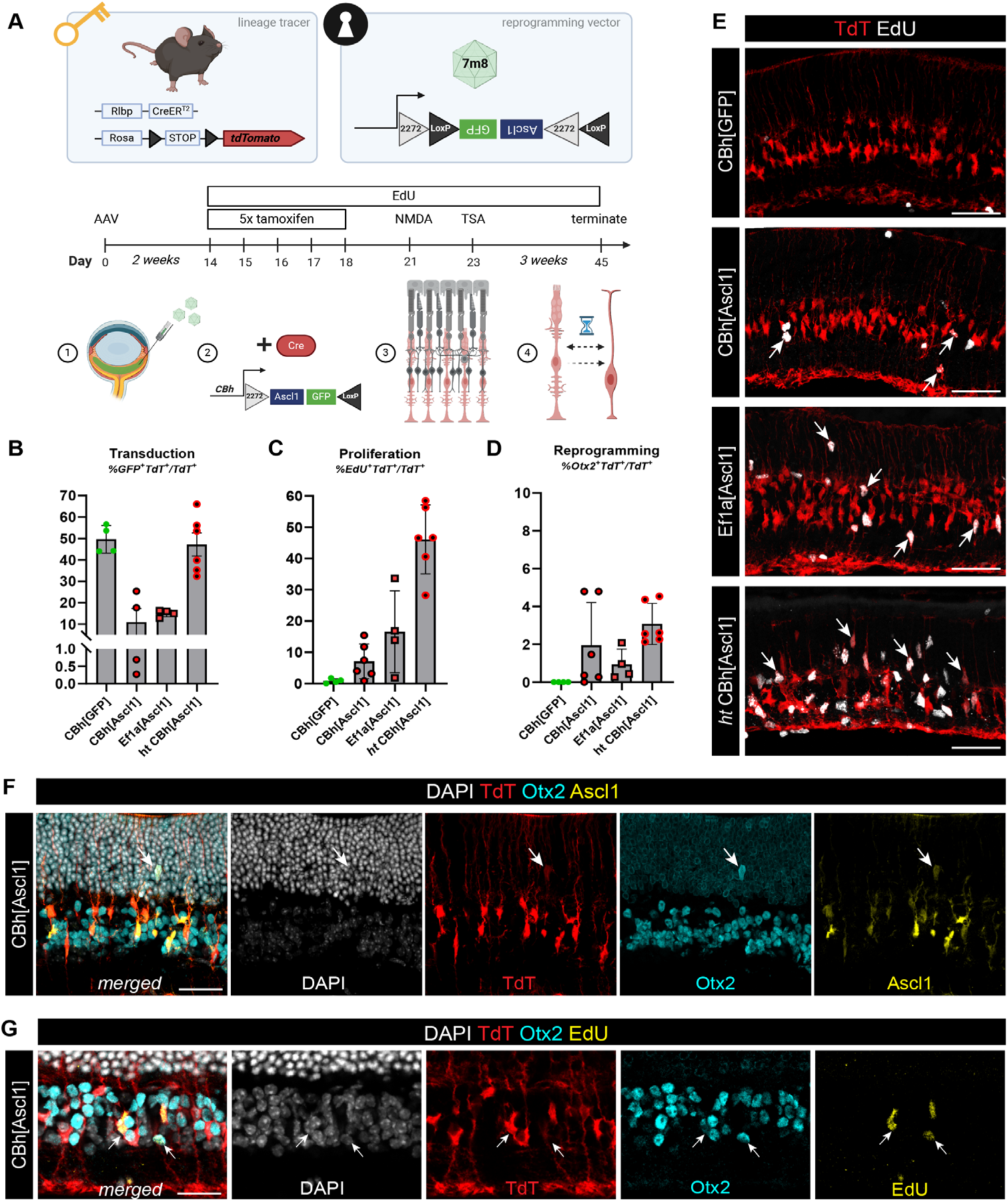
AAV-borne Ascl1 expression in MG stimulate their proliferation and neurogenesis of bipolar neurons. **(A)** schematic of experimental concept and timeline of each intervention in vivo, **(B)** bar plot of transduction efficiency per condition quantified as percentage ratio of GFP+TdT+ cells over all TdT+ with each dot being a biological replicate, **(C)** bar plot of MG proliferation counted as a ratio of EdU+TdT+ over all TdT+ cells with each dot being a biological replicate (error bars: standard deviation), **(D)** bar plot of MG reprogramming counted as a ratio of Otx2+TdT+ over all TdT+ cells with each dot being a biological replicate (error bars: standard deviation), **(E)** fluorescence images of MG proliferation per condition showing EdU in white and TdT in red, with double-labelled examples indicated with white arrows, **(F-G)** fluorescence images of reprogrammed cells (white arrows) per condition showing merged and single channels of DAPI in white, TdT in red, Otx2 in cyan and Ascl1 or EdU in yellow; scalebar for E: 50 μm, for F-G: 20 μm

To effectively lineage-trace MG and their progeny we used a transgenic line where Cre recombinase is expressed under the control of the Retinaldehyde Binding Protein 1 (Rlbp1) promoter and translocates to the nucleus only after exposure to tamoxifen labelling MG with TdT (Rlbp1-CreERt2 x LSL-TdT) (Fig S1A). We have previously reported that this transgenic line restricts Cre and TdT expression to MG ^26,27^ and verified a mean ∼70% recombination efficiency in Sox2^+^ MG after 4-5 tamoxifen injections (Fig S1B) without any detectable co-localization with HuC/D^+^ amacrine/retinal ganglion cells (Fig S1C-C’’) or Otx2^+^ photoreceptors or bipolar neurons (Fig S1D-D”). In the absence of tamoxifen, we observed minimal background signal of TdT in adult animals (2-3 months old), which was confined to Sox2^+^ MG (Fig 1SE-E’, 43 TdT^+^ cells on retinal flat-mount).

To determine the optimal AAV incubation period, we injected adult Rlbp1-CreERt2 x LSL-TdT mice (that were not given tamoxifen) with AAV.7m8/Rlbp1-Cre and assessed TdT reporter expression at 2-, 4-, 6- and 8-weeks post injection (Fig S1F). Vector-treated eyes had detectable TdT signal in Sox2^+^ MG already after 2-weeks incubation, which increased and plateaued after 4-6 weeks (Fig S1G-G’); we therefore concluded a 2-week incubation would be sufficient to obtain AAV-derived expression in the adult mouse retina. After a two-week incubation period of the Ascl1-expressing vectors *in vivo*, mice were given 5 daily doses of tamoxifen to trigger the recombination of the FLEX cassette and label MG with TdT (Fig 1A). We evoked a damage response in the retina by injecting an excitotoxic dose of NMDA intravitreally, followed by an intravitreal dose of TSA (Fig 1A). In our transgenic mouse studies, we allowed the animals to survive for 3 weeks after the injury and TSA; we therefore used a similar survival period for the AAV-derived Ascl1 expression. Three weeks after the NMDA/TSA injections, we sacrificed the animals and assessed the histology of the retina for evidence of AAV transduction (Fig 1B), MG proliferation (Fig 1C) and neurogenesis (Fig 1D).

In the initial round of experiments, we tested two strong ubiquitous promoters, Ef1α and CBh, with the latter at two different titers. Transduction efficiency was assessed by the number of GFP^+^/TdT^+^ cells in three regions across the retinas. The control vector transduced half of TdT^+^ MG (mean=49%, n=4 animals), the low-titer constructs with either CBh or Ef1α promoters driving Ascl1 yielded a modest transduction efficiency (mean=10-15%, n=4 animals), and the high-titer construct with the CBh promoter driving Ascl1 also transduced nearly half of TdT^+^ MG (mean=47%, n=6 animals) (Fig 1B).

We found that all three vectors driving Ascl1 triggered MG proliferation (Fig 1C,E), with patches of TdT^+^ MG expressing EdU and with nuclei either in the inner nuclear layer (INL) or translocated across the apical-basal axis in the outer nuclear layer (ONL) or inner plexiform layer (IPL). These EdU^+^TdT^+^ MG were only found in the retinas treated with Ascl1, and none were present in the retinas of eyes injected with the control vector expressing only the fluorescent reporter (Fig 1E, white arrows). The rare EdU^+^ cells detected in the control condition were not TdT^+^ and most likely microglia, as they localized in the plexiform or nerve fiber layers. As noted, the efficiency of transduction varied across vectors, but irrespective of the range in transduction, all constructs expressing Ascl1 stimulated MG proliferation, with the high-titer vector having the highest mean effect of 46% EdU^+^TdT^+^ MG (n=6 animals)(Fig 1C).

In addition to MG proliferation, vector-borne expression of Ascl1 drove some MG to become Otx2^+^ bipolar neurons (Fig 1D,F,G), similar to what we obtained from previous studies of transgenic reprogramming ^13,16^. The low-titer constructs driving Ascl1 yielded ∼1-2% reprogramming, with the CBh promoter producing a slightly stronger effect than Ef1α (CBh mean=1.9% versus Ef1α mean=0.9%). This efficiency was nearly doubled with the high-titer construct where neurogenesis averaged 3% (Fig 1D). Lineage-traced Otx2^+^ bipolar neurons were often mislocated in the ONL (Fig 1F) and a subset of them co-localized with EdU (Fig 1G), confirming they are *bona fide* newborn neurons.

To further characterize the effects of AAV-borne Ascl1 expression, we used fluorescence-activated cell sorting (FACS) to collect TdT^+^ cells pooled from several infected animals (n≥3) and then processed them for single-cell RNA sequencing (scRNA-seq) (Fig 2A). Any residual cells that were not sequenced were stained confirming we sorted TdT^+^Sox2^+^ MG and TdT^+^GFP^+^Otx2^+^ bipolar cells from control and reprogrammed retinas, respectively (Fig S2A-C). When plotting all the sequenced cells from the experimental conditions in a single integrated graph of reduced UMAP space, we could identify cell clusters based on common transcriptional profiles (Fig 2B). Most of the cells were MG, some of which were reactive (Gfap^+^, Ifit3^+^, Stat1^+^), and smaller clusters were identified as microglia (Ptprc^+^, Tnf^+^, Aif1^+^), astrocytes (Pax2^+^, S100b^+^), rod (Nrl^+^), and cone photoreceptors (Arr3^+^) (Fig 2B), most likely carried over during sorting. We have encountered this caveat of imperfect sorting in our previous experiments of transgenic reprogramming ^13–16^, which is why we verify *bona fide* neurogenesis via histology and EdU co-expression with neuronal markers. Since we do not observe EdU^+^ cells with definitive rod/cone markers, we cannot prove these were newly generated, and so do not include them in the further analyses of MG-derived neurons.

**Fig. 2.**
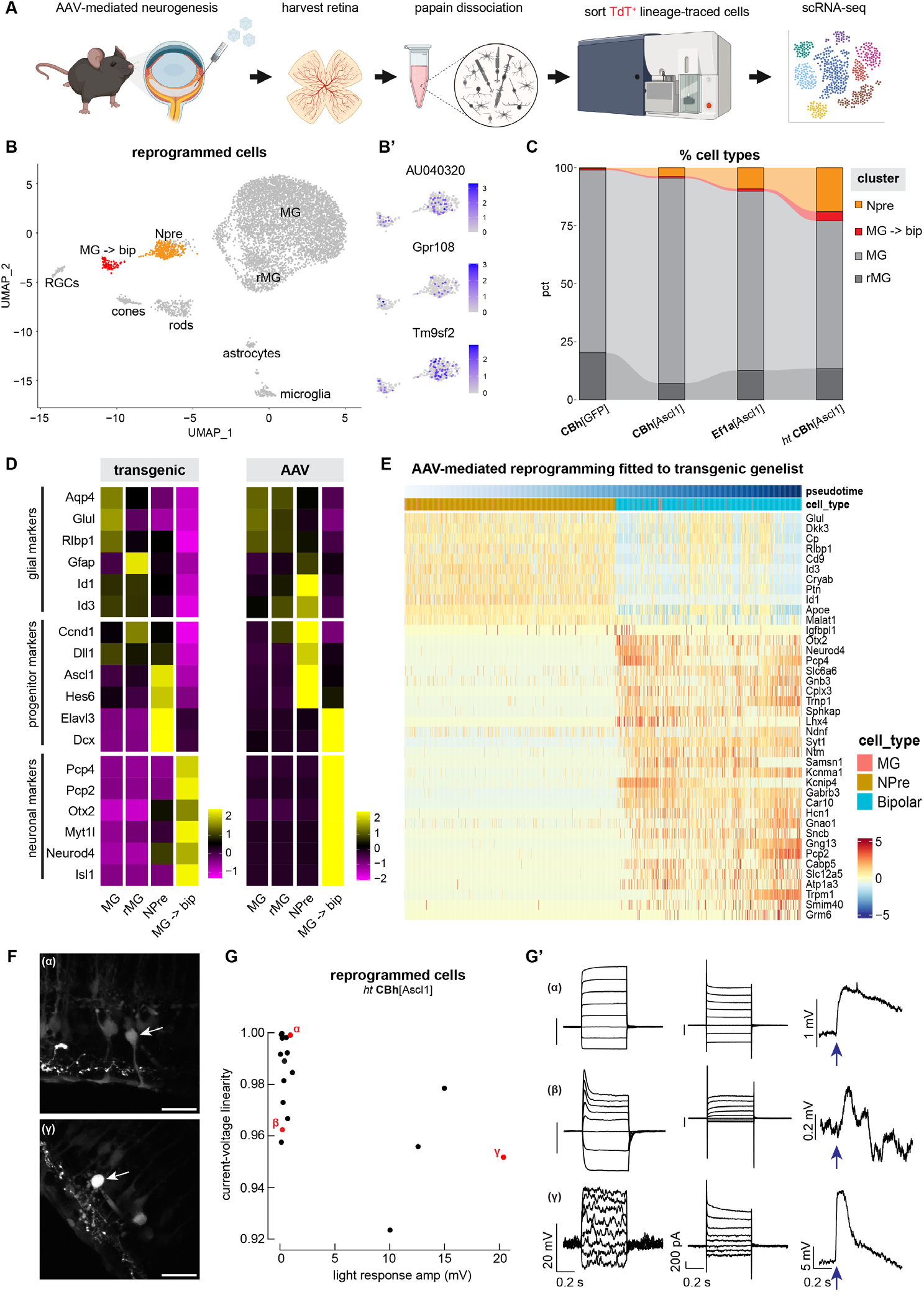
Transcriptional and functional profiling of AAV-mediated reprogramming. **(A)** schematic of experimental pipeline, **(B)** UMAP of integrated scRNAseq data from AAV-borne Ascl1 and control vectors clustered by cell type and highlighting reprogrammed clusters in orange/red, **(B’)** feature plots of subset reprogrammed clusters showing the level of expression for AAV receptor genes, **(C)** alluvium plot of each vector condition showing the proportion of clusters NPre, MG-derived bipolar cells (MG S→ bip), MG and reactive MG (rMG) across samples, **(D)** heatmaps of average gene expression level for select glial, progenitor and neuronal markers in rows and integrated scRNAseq data clusters in columns from published transgenic ^13,16^ or AAV reprogramming experiments, **(E)** heatmap of gene expression across pseudotime of AAV-mediated reprogramming fitted to the top 40 differentially expressed genes identified from transgenic scRNA-seq data plotted across pseudotime as they shift from glial to neurons, **(F)** fluorescent images of representative cells (α and γ) in live retina slices used for patch-clamp recordings, **(G)** scatterplot of linearity values for current-voltage relations for all recorded cells from retina samples reprogrammed with the high-titer AAV/CBh[Ascl1] with red dots indicating example cells α β γ whose profiles are shown in G’, **(G’)** from left to right, family of voltage responses to current steps, family of current responses to voltage steps, and voltage responses to a brief light flash at the time of the arrow.

Of all the cells we analyzed, two cell clusters were distinct, in that they expressed MG transcripts, and also Ascl1 and its downstream targets including Otx2. These cells were labelled as Ascl1^+^ neurogenic precursors (NPre) and MG-de-rived bipolar neurons (MG→bip) (Fig 2B). Both clusters expressed genes AAVR (AU040320), Gpr108 and Tm9sf2 that are reportedly essential for AAV transduction ^28–30^ (Fig 2B’). Cells in these clusters originated mostly from animals treated with Ef1α-

FLEX[Ascl1] and high-titer CBh-FLEX[Ascl1] (Fig 2C, orange and red), which confirmed our histological observations on proliferation and reprogramming (Fig 1C-D). Interestingly, we observed no correlation between vectorborne neurogenesis and MG reactivity, as our control vector (CBh-FLEX[GFP]) accounted for the majority of reactive MG (Fig 2C, dark grey). This is an encouraging indication that transducing MG with AAVs doesn’t evoke an inflammatory response that prevents reprogramming, as we detect both proliferating MG (Fig S2E, TdT^+^EdU^+^ white arrows) and proliferating microglia (Fig S2E, Iba1^+^EdU^+^ yellow arrows) in retinas transduced with CBh-FLEX[Ascl1].

The NPre uniquely expressed Ascl1 and the Notch ligand Delta1 (Dll1), whereas MG-derived neurons expressed the downstream effector Myt1l, neurogenic marker Neurod4 and bipolar marker Otx2 while retaining a low relative expression of the MG marker Rlbp1 in both clusters (Fig S2D). We also detected transcripts of neuronal markers expressed early during neurogenesis and often retained as neurons mature, such as Pcp4, Elavl3, Meis2, Caln1, Isl1, Ebf1, and Pou2f2 (Fig S2D).

The glia-to-neuron conversion we obtained with AAVs phenocopies our results from transgenic animals (Glast-CreERt2 x LSL-tTA x tetO-Ascl1-GFP), even though the glial promoter and mode of Ascl1 expression were different. Specifically, when we integrated the scRNAseq data from either transgenic ^13,16^ or AAV-mediated expression of Ascl1, we confirmed that the pattern of expression for known marker genes was similar between the two (Fig 2D). The average expression of glial genes was highest in MG and reactive MG for both transgenic and AAV-mediated neurogenesis, with STAT-pathway targets Id1/3 being upregulated in NPre of AAV-treated cells (Fig 2D). This aligns with our previous observation that MG failing to become neurons retain high levels of these inhibitor of differentiation genes ^31^ and explains why we see low levels of Rlbp1 expression in the reprogrammed clusters.

Furthermore, we asked how similar the trajectory of reprogramming is between transgenic and AAV-mediated neurogenesis. To address this, we performed pseudotime analysis on an integrated scRNA-seq dataset from previous transgenic experiments (Fig S2F)^13,14,16^ and our new AAV experiments. With pseudotime analysis, where the age/maturity of cells is ranked computationally based on their transcriptomic content, we chose the starting and ending nodes to track the MG-to-neuron conversion and queried which genes were differentially expressed across the trajectory. We obtained a genelist of top 40 genes that followed the trajectory of transgenic MG reprogramming (Fig S2G) and then asked how well the trajectory of AAV-mediated MG reprogramming fit that sequence of gene expression (Fig 2E). Despite the differences in cell number that correspond to each cluster (MG, NPre, MG-derived neurons) between transgenic and AAV-mediated reprogramming, the pattern of gene expression is mostly shared between the two modes of reprogramming. For example, we see consistent downregulation of glial markers Glul and Rlbp1 as cells transition to neuronal fate, as well as a downregulation of the apolipoprotein gene ApoE and long non-coding RNA Malat1 (Fig 2E, S2G, which are primarily expressed by MG in the retina ^32,33^. Interestingly, the NPre during transgenic reprogramming upregulate Igfbpl1 (Fig S2G), which has been associated with axon regeneration ^34^, however this is less prominent during AAV-mediated reprogramming (Fig 2E). Overall, we found a 23% overlap between genelists across pseudotime trajectories, confirming that vector-mediated reprogramming largely follows the transgenic trajectory when shifting between cell states.

To assess whether newborn cells were functional and integrated in the existing retinal network, we performed patch-clamp recordings of single cells in live retina slices (n=3 animals) that were treated with the high-titer CBh-FLEX[Ascl1] vector (Fig 2F, example cells α and γ). Responses varied considerably across recorded cells. Some cells resembled native MG in many responses. For example, cell α in Figure 2G has a near-linear current-voltage relation, as indicated by the similar spacing of the responses to current and voltage steps, and a small and slow light response. These cells (in the upper left corner in Figure 2G) had considerably higher input resistances than native MG (280 ± 80 MOhm vs 40 ± 17 MOhm, mean

± SEM, 10 reprogrammed cells and 3 native MG), indicating a reduction in the K currents that dominate the electrical properties of native MG. The current-voltage relations of some of the reprogrammed cells (e.g. cell β) were shaped strongly by voltage-activated conductances, most likely K currents. Of the 19 GFP^+^TdT^+^ cells we recorded, 8 had significant light responses, 4 that were quite small and 4 that were substantial. Interestingly, the largest light responses here are ∼10x larger than previous reprogramming experiments ^13^.

Consistent with their physiological responses, the reprogrammed clusters were enriched for both sodium and potassium channel genes. Ascl1^+^ NPre expressed potassium channel genes Kcnq4 and Kcnh7 while the MG-derived neurons expressed sodium channel genes Scn1a, Scn2a and potassium channel genes Kcna5, Kcnb2, Kcnd3 and Kcnh5 (Fig S2D). Furthermore, MG-derived neurons expressed genes for AMPA (Gria1, Gria2), metabotropic (Grm1, Grm2, Grm5, Grm6) and NMDA (Grin1, Grin2a) type glutamate receptors, as well as inhibitory ionotropic (Gabra3, Gabrb3) and metabotropic (Gabbr2) GABA receptors (Fig S2D). This suggests that AAV-borne reprogrammed neurons expressed machinery for both excitatory and inhibitory synaptic inputs, in line with their diverse electrophysiological properties.

### Vector-borne atonal factors synergize with Ascl1 to stimulate neurogenesis from adult Müller glia

We have previously shown that the combination of Ascl1 and the related proneural transcription factor Atoh1, was highly efficient in converting MG to a neurogenic state, with up to 80% of the MG generating neurons in the transgenic animals irrespective of injury ^14,16^. In addition, when Atoh1 was added to Ascl1 the MG-derived neurons adopted an RGC-like fate rather than a bipolar fate. To determine whether similar increases in efficiency and changes in neural fate occur with vector-mediated expression of this transcription factor combination, we designed AAV vectors with a FLEX cassette of Ascl1 and Atoh1 or Atoh7 coding sequences driven off a CBh promoter (Fig 3A). These two AAV vectors were intravitreally injected in adult Rlbp1-CreERt2 x LSL-TdT animals following the same experimental pipeline as described above (Fig 1A) and we analyzed the outcome using the same methods.

**Fig. 3.**
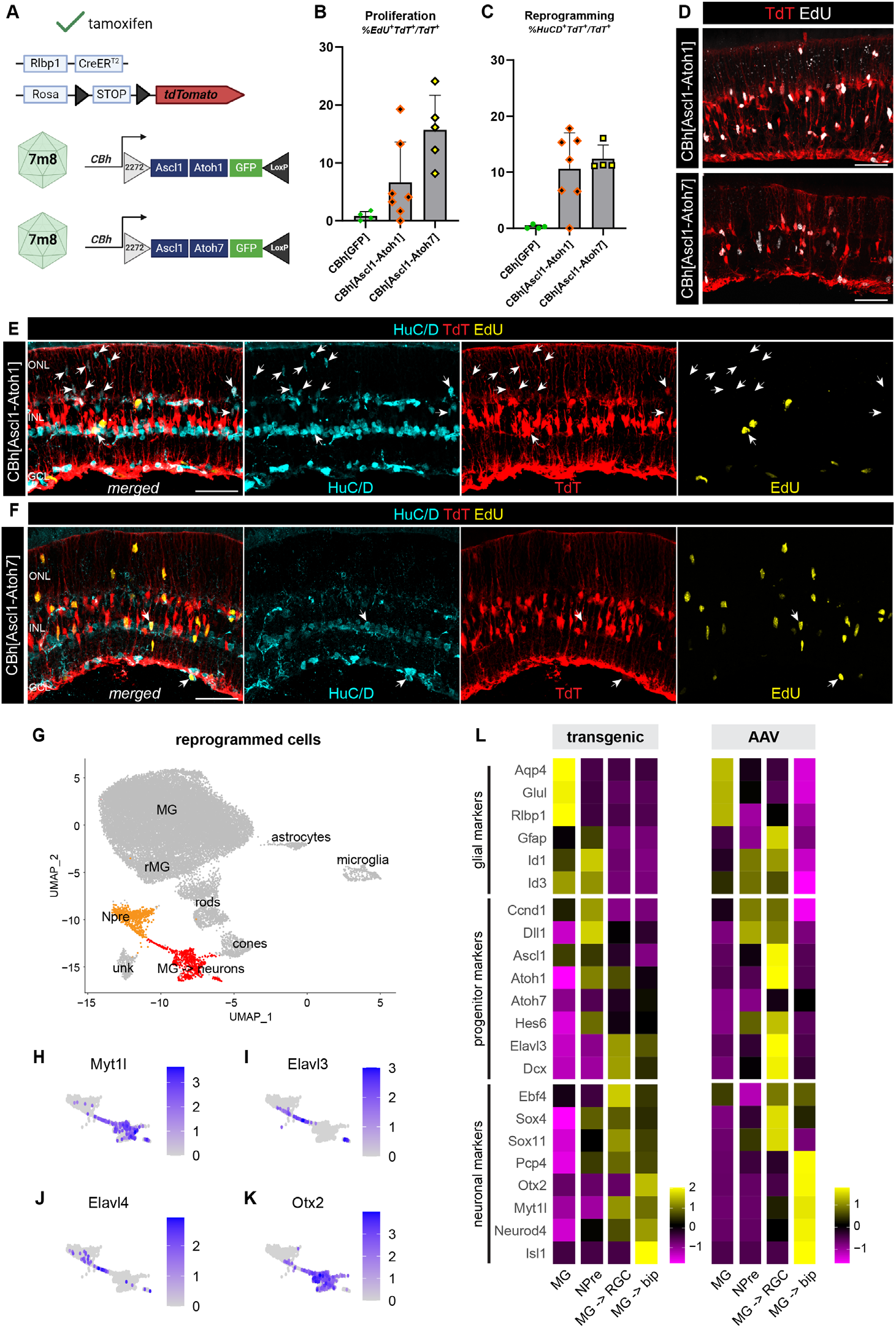
AAV-borne Ascl1-Atoh1/7 expression induces neurogenesis that phenocopies transgenics. **(A)** schematic of AAV vectors and their transgene cassette, **(B)** bar plot of MG proliferation counted as a ratio of EdU^+^TdT^+^ over all TdT^+^ cells with each dot being a biological replicate (error bars: standard deviation), **(C)** bar plot of MG reprogramming counted as a ratio of HuC/D^+^TdT^+^ over all TdT^+^ cells with each dot being a biological replicate (error bars: standard deviation), **(D)** fluorescence images of MG proliferation per condition showing EdU in white and TdT in red, **(E-F)** fluorescence images of reprogrammed cells (white arrows) per condition showing merged and single channels of HuC/D in cyan, TdT in red and EdU in yellow, **(G)** UMAP of integrated scRNAseq data from AAV-borne Ascl1-Atoh1/7 clustered by cell type and highlighting reprogrammed clusters in orange/red, **(H-K)** feature plots of subset reprogrammed clusters showing the level of expression for the gene annotated above each plot, **(L)** heatmaps of average gene expression level for select glial, progenitor and neuronal markers in rows and integrated scRNAseq data clusters in columns from published transgenic or AAV reprogramming experiments; scalebars for D-F: 50 μm, ONL: outer nuclear layer, INL: inner nuclear layer, GCL: ganglion cell layer.

AAV-mediated expression of Ascl1-Atoh1 in adult MG reproduced our observations from transgenic animals ^14^; we found an increase in MG proliferation and detected HuC/D^+^TdT^+^ MG-derived RGC-like neurons. Since our vector payload was customizable and we were not limited by the availability of transgenic mice, we compared the combination Ascl1-Atoh1 (n=7 animals) to Ascl1-Atoh7 (n=5 animals), which we previously reported to evoke neurogenesis from MG *in vitro* but had not yet tested *in vivo* (see figure 1 of Todd *et al*., 2021).

Both combinations were able to induce robust MG proliferation (Fig 3B) and neurogenesis *in vivo* (Fig 3C), with ∼5-15% lineage traced MG incorporating EdU, similar to Ascl1 alone for the low-titer vector (Fig 1C) and showing nuclear translocation across the baso-apical axis (Fig 3D). For both vectors, driving Ascl1-Atoh1 or Ascl1-Atoh7, the main neuronal class obtained were HuC/D^+^ amacrine/RGC-like cells (Fig 3E-F), with some Otx2^+^ bipolar cells generated as well (Fig S3A-B), in line with our previous observations from transgenic reprogramming.

Interestingly, Ascl1-Atoh7 was more efficient at driving the genesis of bipolar cells compared to Ascl1-Atoh1 (Fig S3B). Approximately 10% of lineage-traced MG gave rise to HuC/D^+^ neurons after Ascl1-Atoh1/7 expression (Fig 3C, E, F, white arrows). Some of these cells were ectopically found in the ONL (Fig 3E, white arrows), as we have seen in transgenic animals ^14^, and some incorporated EdU, thus confirming the cells were generated from proliferating progenitors (Fig 3E-F).

Transcriptional profiling of TdT^+^ sorted cells after AAV-mediated reprogramming confirmed our histological observations; we obtained distinct cell clusters of NPre and MG-derived neurons (MG→ neurons) (Fig 3G). These clusters expressed Ascl1 but low levels of Atoh1/7 and TurboGFP, potentially due to the order of genes on the polycistronic vector cassette or transcript downregulation. Importantly, we detected downstream effector markers such as Myt1l (Fig 3H) and neuronal markers Elavl3/4 for amacrine/RGC-like neurons (Fig 3I-J) and Otx2 for bipolar neurons (Fig 3K).

Vector-induced neurons phenocopy the fate and expression profile of glia-derived neurons from transgenic animals ^14,16^, with known genes similarly patterned across MG, NPre, MG-derived RGCs (MG → RGCs) and MG-derived bipolar cells (MG → bip) (Fig 3L). MG-derived RGCs from transgenic animals seem to downregulate glial and progenitor genes more effectively compared to AAV, as the latter still express Gfap and Id1/3. However, for both conditions, MG-derived RGCs express Elavl3, Dcx, Ebf4, Sox4 and Sox11 (Fig 3L), which are genes we previously identified in MG-derived RGCs following transgenic expression of Ascl1-Atoh1^14^ and Islet1-Pou4f2-Ascl1^15^.

Our ability to stimulate neurogenesis using AAVs was most influenced by the vector titer and the promoter, though it was clear that the combinations Ascl1-Atoh1 and Ascl1-Atoh7 had a stronger effect than Ascl1 alone when delivered at the same titer (∼2% for Ascl1, ∼10-15% for Ascl1-Atoh1/7). This improvement in neurogenic potential was true also in our transgenic animals, with Ascl1-Atoh1 converting ∼80% MG into neurons whereas Ascl1 alone would convert ∼30% ^14^. Interestingly when comparing vector-mediated expression of Ascl1-Atoh1 and Ascl1-Atoh7, the latter seems more effective at inducing NPre and new neurons in our experimental context (Fig S3C).

We also performed pseudotime analysis to ask how similarly cells change their fate when the transcription factors are delivered via AAV as opposed to transgenics. To this end, we consolidated previous scRNA-seq datasets from transgenic experiments ^14,16^ and obtained a genelist of top 40 genes that followed the trajectory of transgenic MG reprogramming (Fig S3D). We then asked how well the trajectory of AAV-mediated neurogenesis fit that expression pattern. As we saw for Ascl1, the pattern of gene expression is mostly shared between the two modes of reprogramming (Fig S3D). One obvious difference was the curved expression pattern of Atoh1 and Pcp4 in transgenic reprogramming, where they are first downregulated as the cells are in a glial state, then upregulated as they become progenitors and RGC-like cells and again downregulated as they become bipolar cells. AAV-induced neurons do not follow the same pattern for these genes, as Atoh1 levels seem consistent across states and Pcp4 being upregulated in the glia-derived bipolar cells. As such, vector-mediated reprogramming is not identical to transgenic animals, however overall, there is a similar trajectory of induced neurogenesis *in vivo*.

### AAV-mediated reprogramming is affected by vector incubation period

The main rate-limiting step for AAV-genome expression is second-strand DNA synthesis to generate a double-stranded DNA molecule from which transcription can be initiated ^35^. As such, we hypothesized that a longer incubation period after AAV delivery in the retina could improve the expression of the vector cargo and by extension improve neurogenesis. To test this, we expanded our previous experimental pipeline and waited 6-weeks after AAV injection (Fig 4A-A’) to match the previously reported transduction peak for capsid AAV.7m8 ^25,36^. We used this expanded pipeline of AAV-borne Ascl1 or Ascl1-Atoh1 expression in transduced MG, followed by NMDA injury and TSA, and TdT^+^ cells were subsequently sorted from retinal tissues 3-4 weeks post TSA (Fig 4A’).

**Fig. 4.**
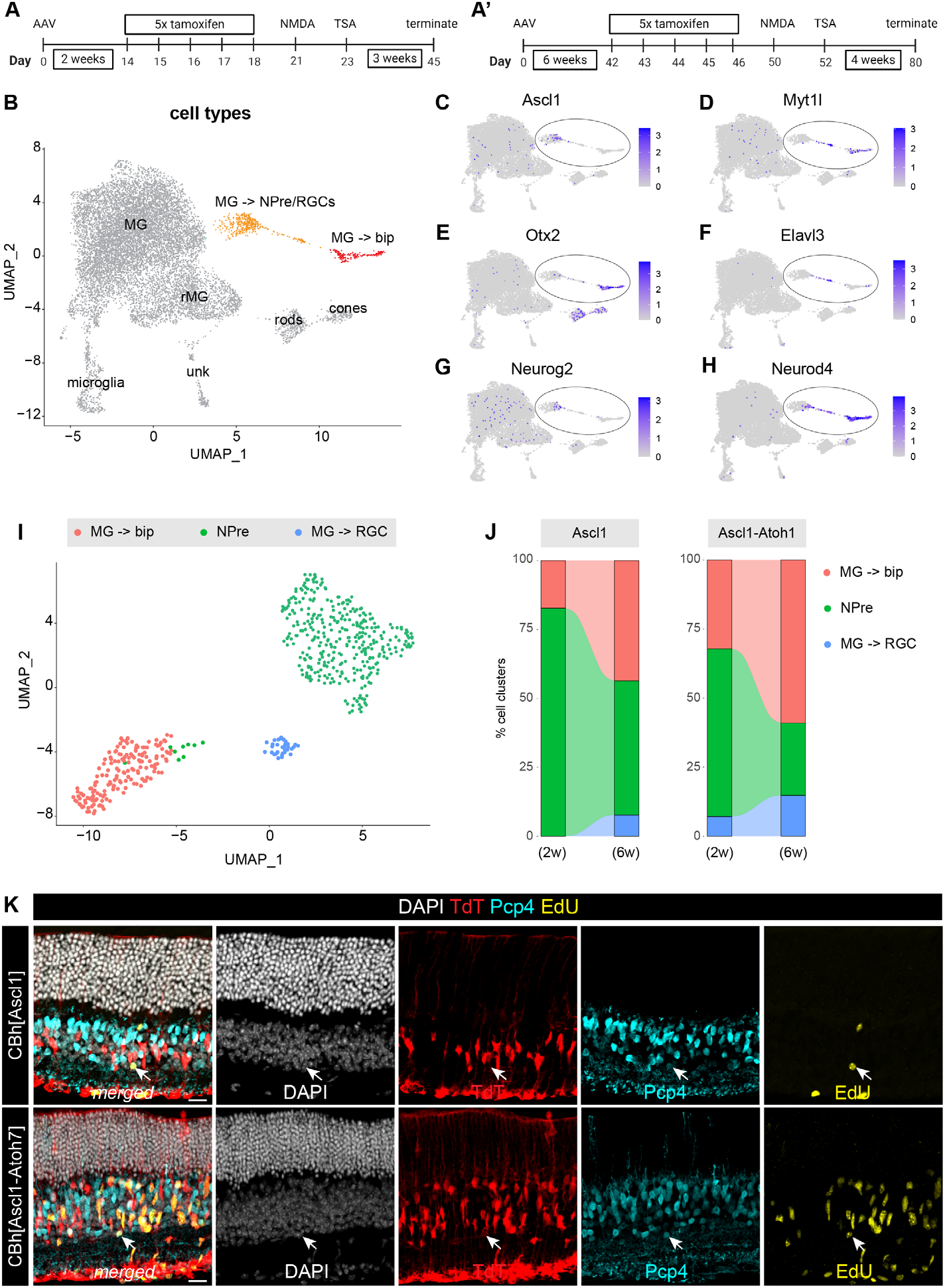
The impact of AAV incubation time on neurogenesis. **(A-A’)** schematic of experimental timeline where AAV is incubated for 2- or 6-weeks, **(B)** UMAP of integrated scRNAseq data from AAV-reprogramming experiments from all timepoints and vectors clustered by cell type, **(C-H)** feature plots of integrated UMAP from (A) showing the level of expression for the gene annotated above each plot with reprogrammed clusters circled, **(I)** reclustered UMAP of reprogrammed cells after integrating all AAV-borne reprogrammed cells for both timepoints, **(J)** alluvium plots of each vector-derived cargo (Ascl1 or Ascl1-Atoh1) showing the percentage of each cell cluster in samples of 2-or 6-week AAV incubation from data subset shown in (I), **(K)** fluorescence images of reprogrammed cells (white arrows) per condition showing merged and single channels of DAPI in white, TdT in red, Pcp4 in cyan and EdU in yellow; scalebar: 20 μm

We integrated the scRNAseq datasets from all conditions, both for short (2-weeks) and long (6-weeks) incubation periods, and defined cell clusters based on their transcript profiles (Fig 4B). The expression pattern of key genes Ascl1, Myt1l, Otx2, Elavl3, Neurog2 and Neurod4 (Fig 4C-H) highlighted the reprogrammed cells (Fig 4B, orange and red clusters). We then subset and re-clustered these cells to specifically compare their distribution across timepoints, and profiled their three main states: cycling NPre, MG-derived bipolar cells and MG-derived RGCs (Fig 4I).

We found that the incubation period of the virus somewhat improved the reprogramming efficiency. The 6-week incubation time samples, for both Ascl1 and Ascl1-Atoh1, showed an increase in the percentage of neurons (either RGC or bipolar) and a relative reduction in the percentage of NPre cells when compared to the 2-week incubation time (Fig 4J). This result is in line with the previously reported kinetics of the AAV.7m8 capsid ^25^ and our own AAV kinetics assay that reproduced the reported plateau of transduction after 6-weeks *in vivo* (Fig S1F-G). We did not observe any evidence that the neurons obtained following longer AAV incubation had a more mature profile, as we were able to obtain mature ON-bipolar neurons (Pcp4^+^TdT^+^EdU^+^) from both Ascl1 and Ascl1-Atoh7 vector-treated eyes after only 2 weeks of AAV incubation (Fig 4K, white arrows). As such, a longer AAV incubation relatively improved reprogramming efficiency but did not affect the maturity of glia-derived neurons.

## Discussion

The ability to stimulate neurogenesis from MG with transient expression of transcription factors provides a potential path to restoring vision in people that have lost retinal neurons due to injury or disease. Up to now, we have relied on transgenic expression of proneural factors, individually or in combinations, to study this approach *in vivo*. These mouse models were instrumental to our understanding of what neuronal classes we can obtain after expressing Ascl1 alone (bipolar cells) or in combination with Atoh1, Islet1 and Pou4f2 (bipolar cells, amacrine/RGC cells) ^13–15^. We were able to decipher important prerequisites for translation, using transgenic animals, such as the importance of retinal injury type and timing relative to proneural factor expression ^16^. To our advantage, MG expressing Ascl1 or Ascl1-Atoh1 can give rise to new neurons following both inner and outer retinal damage, after the neurons have already died ^16^; a necessary condition for treating most retinal diseases.

To generate a regenerative strategy that can someday be used to treat human retinal damage, one approach is to introduce reprogramming factors to MG via AAV vectors. AAV vectors are small icosahedral capsids (∼25 nm) that encapsulate a single-stranded (∼4.7 kb) or double-stranded (∼2.4 kb) DNA cassette flanked by inverted terminal repeats (ITRs). This cassette encodes the information for vector packaging and facilitates intermolecular recombination to form circularized concatemers for episomal persistence of AAV genomes in the host cell nucleus ^37,38^. AAV vectors are currently used in several clinical trials in the eye for gene replacement or gene editing and can drive long-term expression of AAV-payload after just a single administration ^17,18^.

Although AAV-mediated gene delivery has gained in popularity for ocular indications, there are several questions that need to be answered with regards to their use for MG reprogramming as a strategy for retinal regeneration: [1] can AAV-derived expression of proneural genes induce MG proliferation and neurogenesis, [2] can we restrict AAV vector expression to MG specifically, [3]what are the kinetics of AAV expression and does it matter, [4] will AAV vectors trigger a gliotic response that blocks reprogramming? In this report we have tested these features and found that AAV-mediated reprogramming of MG is both traceable and effective.

Our results show that [1] driving transcription factor expression in MG, where tamoxifen-inducible Cre expression is under the control of a transgenic Rlbp1 promoter, can result in high levels of recombination in the adult mouse retina and is sufficient for AAV-mediated expression of FLEX cassettes driven by ubiquitous promoters (CBh and Ef1α). This results in a robust effect of MG proliferation only when Ascl1, Ascl1-Atoh1 or Ascl1-Atoh7 are expressed, but not when a fluorescent reporter is expressed alone. Importantly, vector-mediated expression of these proneural genes led to neurogenesis towards a bipolar and amacrine/RGC-like fate following a similar gene trajectory as we previously reported in experiments with transgenic animals, even though the glial promoters in each case were different, namely Rlbp1 and Glast. The neurons we obtained after AAV-mediated reprogramming were integrated in the retinal network, had electrophysiological responses to light and expressed genes of the machinery required for both excitatory and inhibitory synaptic inputs. Strikingly, the maximum light response amplitude we obtained from several of these cells was much larger than reprogrammed cells from our previous transgenic experiments ^13,15^ and approached amplitudes characteristics of endogenous neurons. Thus AAV-mediated reprogramming produced cells that effectively integrated into retinal circuits. Electrically, however, these cells differed less from MG than those we characterized in transgenic experiments; specifically, voltage-activated conductances in cells from AAV-mediated reprogramming were considerably smaller than those from transgenic experiments.

With regards to specificity, recent reports have claimed AAV-mediated neurogenesis when in fact there was ectopic expression of vector payload in endogenous neurons or insufficient controls of vector specificity ^23,24,39^. This has been addressed by follow-up studies that demonstrated the artefacts and highlighted caveats of using AAVs to induce cell fate changes, both in the brain and retina ^20,22^. One of the causes for ectopic AAV-cargo expression was the use of the glial-specific GFAP mini promoter, used to drive expression in astrocytes and MG ^23,24,39^. The specificity of the GFAP promoter was carefully examined and shown to leak depending on the genes downstream, especially when driving proneural transcription factors such as NeuroD1, Ascl1 and Atoh7 ^21^. In the same report however, FLEX cassettes under the control of a ubiquitous Ef1α promoter that drove proneural genes to MG did not result in leaky expression of AAV cargo in endogenous neurons, but also showed no evidence of induced neurogenesis from MG ^21^. In our study we show that [2] restricting AAV-payload expression in MG can be achieved when the specificity originates from a transgenic Rlbp1 promoter. Importantly, [3] this specificity does not change over longer periods of AAV incubation (2-8 weeks), even when the Rlbp1 promoter is delivered via AAV. In fact, adapting the experimental conditions to the optimal AAV kinetics (from 2 weeks to 6 weeks incubation) can slightly improve reprogramming efficiency.

We have shown that delivering Ascl1 or Ascl1-Atoh1/7 to MG with AAVs, and controlling their expression temporally with the FLEX system, can drive glia-to-neuron conversion *in vivo* and produce EdU^+^ bipolar, amacrine/RGC-neurons analogous to previous transgenic experiments. The reactive gliosis evoked by the AAV injection and transient transgene expression, as well as the intravitreal NMDA and TSA injections, did not prevent MG from changing fate with our paradigm. We show that [4] the reactivity of MG was comparable across vector treatments, even at a high AAV titer, and similar in gene profile to the equivalent cluster from transgenic experiments. In large animal studies on non-human primates and human clinical trials, immunosuppression is administered prior to intraocular AAV injections as standard practice ^6,40^, which we envision would also become part of a future AAV-based regenerative regimen.

This body of work serves as a bridge towards our ultimate goal of an all-encompassing AAV strategy to stimulate neurogenesis by expressing proneural factors in MG of adult patients. We dissected the issues of traceability and efficiency into a transgenic and vector component respectively, to demonstrate that AAV-borne expression of proneural factors is capable of altering cell fate in the adult mammalian retina. Now the challenge is to improve the efficiency of neurogenesis and design a strategy that is solely based on AAV-derived genetic material. By incorporating lessons learnt from previous studies and the ongoing research on AAV biology, we can devise a reliable strategy that stimulates neurogenesis from MG using one or multiple viral vectors ^41^. Such a regimen would have immense therapeutic impact and could be applied in a disease-agnostic manner. We believe this work makes a significant contribution to achieving this goal as it proves we are able to generate new neurons in an adult mammalian retina using vector-delivered cargo.

## Acknowledgements

We would like to thank all the members of the Reh lab and the Bermingham-McDonogh lab for their valuable comments on the manuscript. All schematics were generated using Biorender. This paper was typeset with the bioRxiv word template by @Chrelli: www.github.com/chrelli/bioRxiv-word-template

## Author contributions

Conceptualization: MP, TAR

Methodology: MP, MPr

Investigation: MP, MPr, EF, LK, ARP, FR

Visualization: MP, MPr, EF, FR

Supervision: TAR

Writing—original draft: MP, TAR

Writing—review & editing: MP, TAR

## Competing interest statement

Some of the findings in this report are part of a patent application that has been submitted by the University of Washington: Patent Application 63/362,361 filed 4 January 2022. TAR is a co-founder of Tenpoint Therapeutics Ltd. All other authors declare they have no competing interests.

## Data and materials availability

All data are available in the main text or the supplementary materials. Some of the scRNAseq analysis was done using previously published data and are referenced accordingly. All sequencing data will be deposited to an online database (GEO pending).

## Materials and Methods

### Animals

All animals were treated and housed with University of Washington Institutional Animal Care and Use Committee approved protocols. Males and females were both used in experiments at equal frequencies. All experiments were performed on adult mice that were over 30 days old. For EdU incorporation, animals were given 0.4 mg/ml EdU/H2O ad libitum from the first day of intraperitoneal tamoxifen injections onwards until the animals were terminated.

### Injections

Intravitreal injections were performed with a 32-G Hamilton syringe on mice anesthetized with isoflurane. AAV preps were produced by Vectorbuilder. AAV injections were done in a volume of 1.5-2 μl and the vector titers are outlined in the table below. Injections of NMDA were done in a volume of 1 μl at a concentration of 100 mM in PBS. TSA (Sigma-Aldrich) was administered via intravitreal injections in DMSO at a concentration of 1 μg/μl. Intraperitoneal injections of tamoxifen (1.5 mg per 100 μl of corn oil) were administered to adult mice for five consecutive days to induce recombination of the AAV.7m8/FLEX cassette.

**Table 1.** AAV vectors and their respective titers.

| Vector cassette | Titer GC/ml |
| --- | --- |
| CBh>FLEX[EGFP]-WPPE | 2.15x10<sup>12</sup> |
| CBh>FLEX[hASCL1(ns):T2A:TurboGFP]-WPPE | 3.93x10<sup>11</sup> |
| CBh>FLEX[hASCL1(ns):T2A:TurboGFP]-WPPE | 4.60x10<sup>13</sup> |
| Ef1α>FLEX[hASCL1(ns):T2A:TurboGFP]-WPPE | 8.44x10<sup>11</sup> |
| CBh>FLEX[hASCL1(ns):hAHOH7(ns):P2A:TurboGFP]-oPRE | 7.92x10<sup>11</sup> |
| CBh>FLEX[hASCL1(ns):T2A:hAHOH1(ns):P2A:TurboGFP]-oPRE | 5.62x10<sup>11</sup> |

### Immunohistochemistry

Dissected eye cups were fixed with 4% paraformaldehyde/PBS for 30 minutes and then incubated in 30% sucrose solution at 4ºC overnight. Eyes were then embedded in optimal temperature cutting compound before freezing. Frozen samples were sectioned at -20 ºC in 15-to 18-µM sections onto glass slides. Slides were then heated for 10 min on a slide warmer before staining or freezing at -20 ºC for long-term storage. For staining, slides were traced with a liquid blocker pen and then rehydrated with PBS. For EdU detection the Click-iT™ EdU Imaging Kit (Thermo Fisher Cat # C10086) was used following manufacturer instructions, followed by antibody staining. Primary antibodies were incubated overnight at 4ºC in blocking solution (0.5% TritonX-100 and 5% Normal Horse Serum in PBS). The primary solution was removed, and slides were washed with PBS. Secondary antibodies were incubated in blocking solution for 90 min and then slides were washed with PBS. Fluoromount-G (Southern-Biotech) mounting medium was added to slides before covering slide with a glass coverslip. See table below for all antibodies used.

**Table 2.**
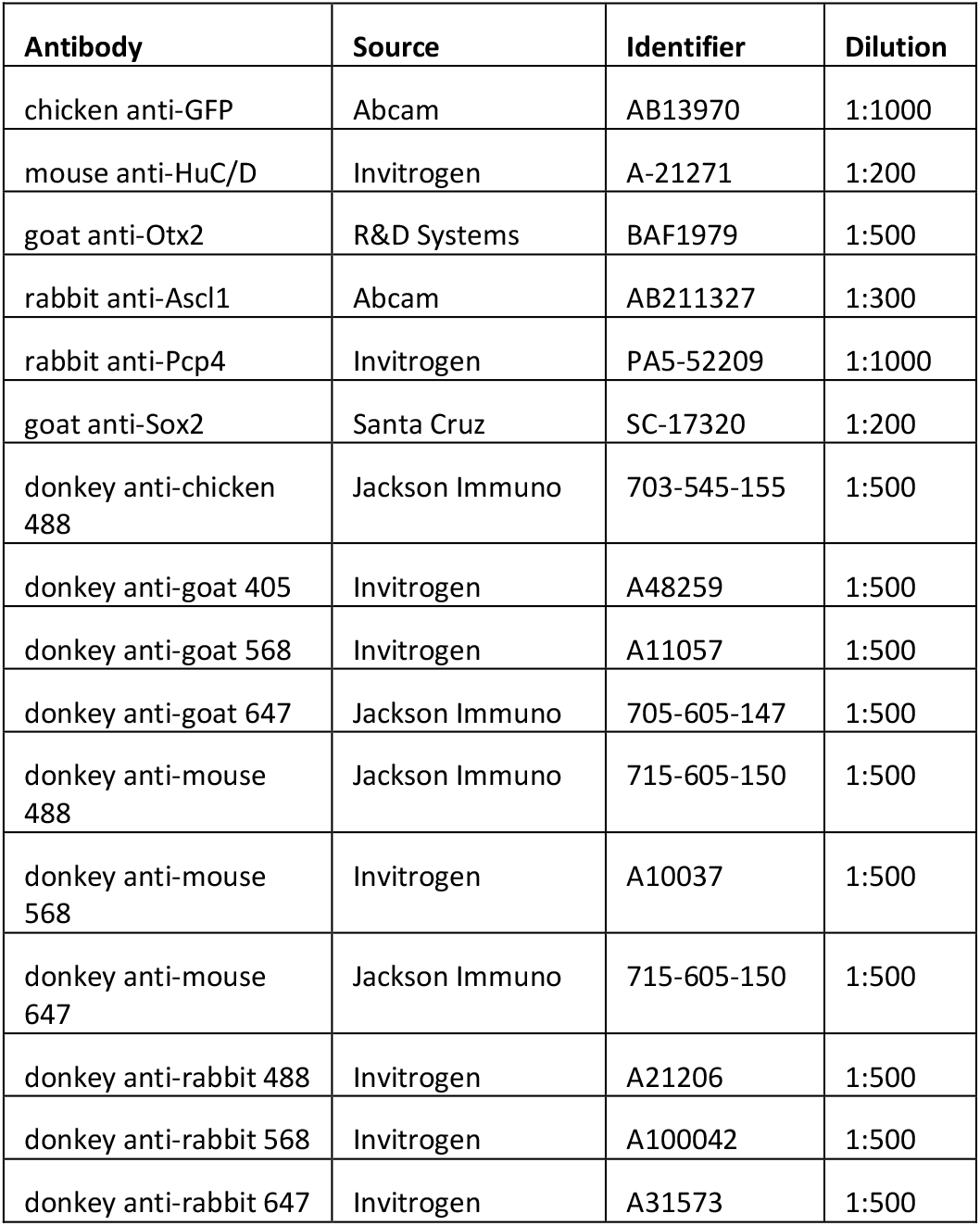
List of primary and secondary antibodies.

### Microscopy

Stained sections were imaged using a Zeiss LSM800 microscope. For quantification, images were taken with a 20x objective with at least 3 images taken per retina. Images were analyzed and counted using FIJI. Lineage-traced TdT+ cells were initially counted from maximum intensity z-projections. The colocalization of markers was then identified and verified by going through single-planes of the z-stack. Careful attention was paid to ensure there was a consistent pattern of marker overlap in each plane where the cell was present.

### Fluorescence-activated cell sorting

Following euthanasia, pools of four retinas were dissociated using the Worthington Papain Dissociation System (catalog no. LK003150) according to the manufacturer’s instructions. Cells were then spun at 4 °C at 400 × g for 10 min and resuspended in neurobasal medium (Gibco no. 21103049), 10% fetal bovine serum (Clontech), B27 (Invitrogen), N2 (Invitrogen), 1 mM l-glutamine (Invitrogen), and 1% penicillin-streptomycin (Invitrogen). The cell suspension was passed through a 35-μm filter and then sorted using a BD FACSAria III cell sorter (BD Bioscience) to retrieve TdT+ cells. During sorting, appropriate gates were implemented to exclude debris and doublets. A minimum of 40,000 events were collected in tubes that were coated with 10% BSA.

### Electrophysiology

Recordings were performed identical to our previous reports ^15^. Mice were dark-adapted before recordings. After euthanasia, retinas were sliced into 200-μm slices for recording. Tissue recordings were performed in Ames medium at 32°C and oxygenated with 95% O_2_/5% CO_2_. GFP^+^ cells were targeted for recording using video differential interference contrast with infrared light and confocal microscopy. Light responses were measured under infrared conditions, and the tissue was exposed to full-field illumination via a green light-emitting diode. Recordings were performed using pulled glass pipettes and filled with solution containing the following: 123 mM K-aspartate, 10 mM Hepes, 1 mM MgCl2, 10 mM KCl, 1 mM CaCl_2_, 2 mM EGTA, 0.5 mM tris–guanosine triphosphate, 4 mM MG–adenosine triphosphate, and 0.1 mM Alexa Fluor 695 hydrazide. Current-voltage relations were measured by making steps to a series of voltages and measuring the resulting currents; linearity of these current-voltage relations was measured by the R^2^ value of a straight line fit to the data.

### Single cell RNA library construction

FACS-purified TdT^+^ MG cells were spun at 4 °C at 400 × g for 10 min and resuspended at a concentration of 1,000 cells/μL. Library construction was performed using the Chromium Next GEM Single Cell 3′ version 3.1 (dual index) protocol and reagents according to the manufacturer’s instructions.

### Since cell RNA sequencing, mapping and data analysis

Multiplexed libraries were sequenced using an Illumina NextSeq 500 using high-output 150 kits. Data were demultiplexed and aligned to the mouse mm10 genome using Cell Ranger Count version 7.2 ^42^. Filtered output files were further analyzed in R using Seurat version 4.3.2 ^43^, ggplot2, data.table, dplyr, tidyr, and other commonly used R packages. Low-quality cells (identified as having low read depth or high mitochondrial content; >10%) were removed from datasets. Before analysis, the cell number was downsampled to the object with the lowest cell number and cell unique molecular identifiers were downsampled to the lowest median for each object using the package scuttle’s downsampleMatrix function. Cells were clustered using principal-components analysis and UMAP. Comparisons between datasets were made by canonical correlation analysis, as described by the Satija laboratory vignette (https://satijalab.org/seurat). Alluvial bar plots were created by the percentage of cells in each cluster split by each sample using ggplot2 with ggalluvial. Heatmaps was calculated using Seurat’s DoHeatmap over the top differentially expressed (DE) genes or selected genes for each cell type cluster with their average expression of that gene. The top DE genes were found by using FindAllMarkers and sorted by the average log_2-fold change for each single cluster compared against all others. Pseudotime-based trajectory analysis was conducted on the integrated RDS files using Monocle3 according to the Trap-nell laboratory vignette (https://cole-trapnell-lab.github.io/monocle3/). The Seurat objects were converted to Monocle3 objects using ‘as.cell_data_set()’. Cells were reclustered using ‘cluster_cells()’, and the principal graphs were calculated using ‘learn_graph()’. Cells were ordered in pseudotime using ‘order_cells()’ with the start nodes chosen based on expressions of marker features. Paths on the graphs were selected using ‘choose_graph_segments()’ by specifying the start and end nodes. The paths were subset using Seurat’s ‘subset()’. The cells were reclustered, and the graphs were relearned in Monocle3. Gene lists of the top 40 differentially expressed (DE) genes were identified based on the highest Moran’s I values obtained from ‘graph_test()’. Expression of the top DE genes was plotted using ComplexHeatmap’s ‘Heatmap()’.

**Figure S1:**
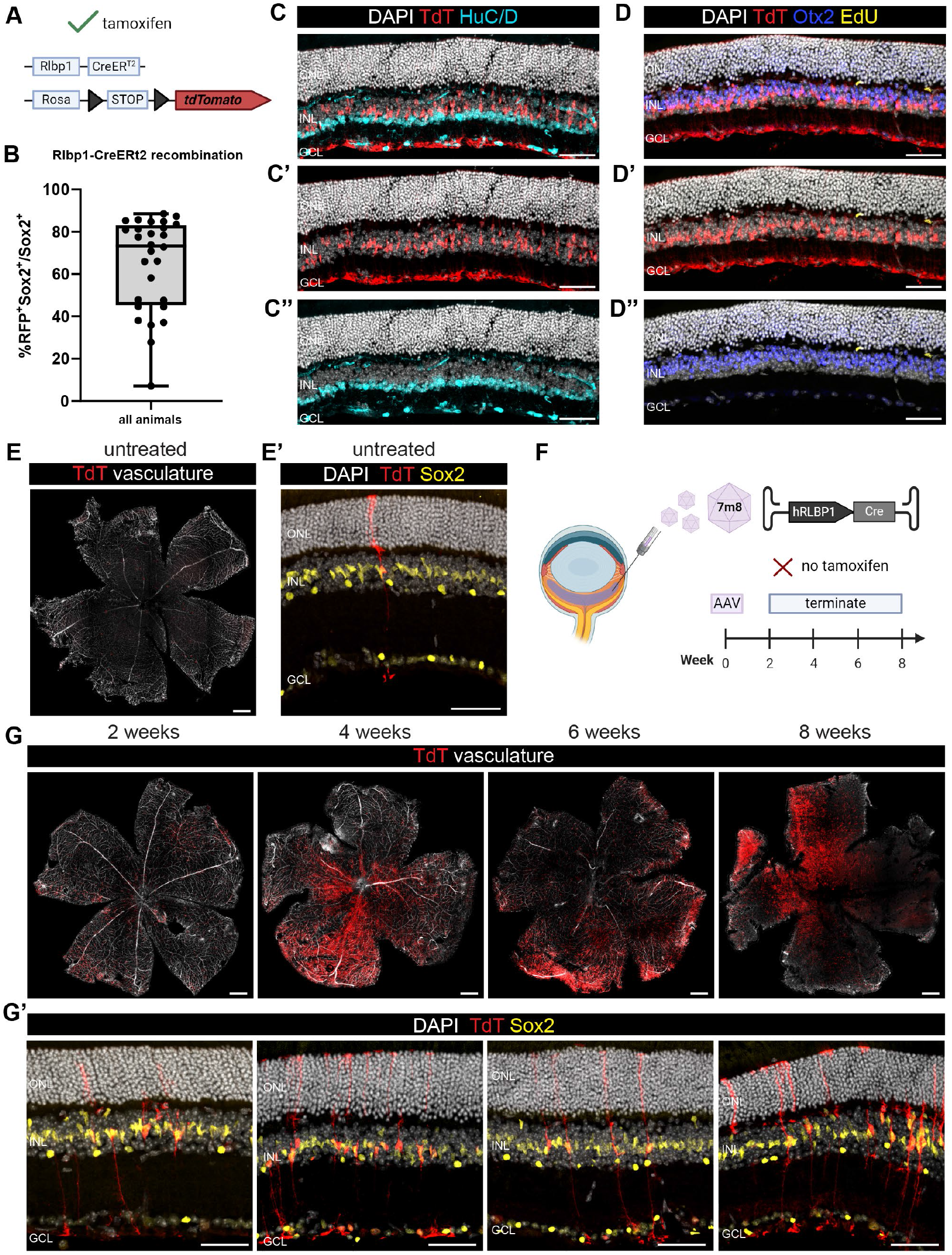
Assessment of transgenic lineage tracer line and AAV kinetics *in vivo*. **(A)** schematic of tamoxifen-inducible transgenic mouse line for MG-specific expression, **(B)** bar plot quantification of recombination efficiency counted as percentage ratio of RFP+Sox2+ cells over all Sox2+ where each dot is a biological replicate (error bar: standard deviation), **(C-C’’)** fluorescence images of Rlbp1-CreERt2 x LSL-TdT retina cross-sections after tamoxifen showing DAPI-stained nuclei in white, TdT in red and HuC/D in cyan, **(D-D’’)** fluorescence images of Rlbp1-CreERt2 x LSL-TdT retina cross-sections after tamoxifen showing DAPI-stained nuclei in white, TdT in red, Otx2 in blue and EdU in yellow, (E) fluorescence image of an untreated retina flatmount with TdT in red and vasculature in white, (E’) representative retina cross-section of untreated retina with TdT in red and Sox2 in yellow (F) schematic of experimental design, (G) fluorescence images of retina flatmounts for each timepoint with TdT in red and vasculature in white, (G’) representative retina cross-sections from timepoints in (G) with DAPI-stained nuclei in white, TdT in red and Sox2 in yellow; scalebars for C-C’’, D-D’’, G’, E’: 50 μm, for E, G: 500 μm, ONL: outer nuclear layer, INL: inner nuclear layer, GCL: ganglion cell layer

**Fig. S2.**
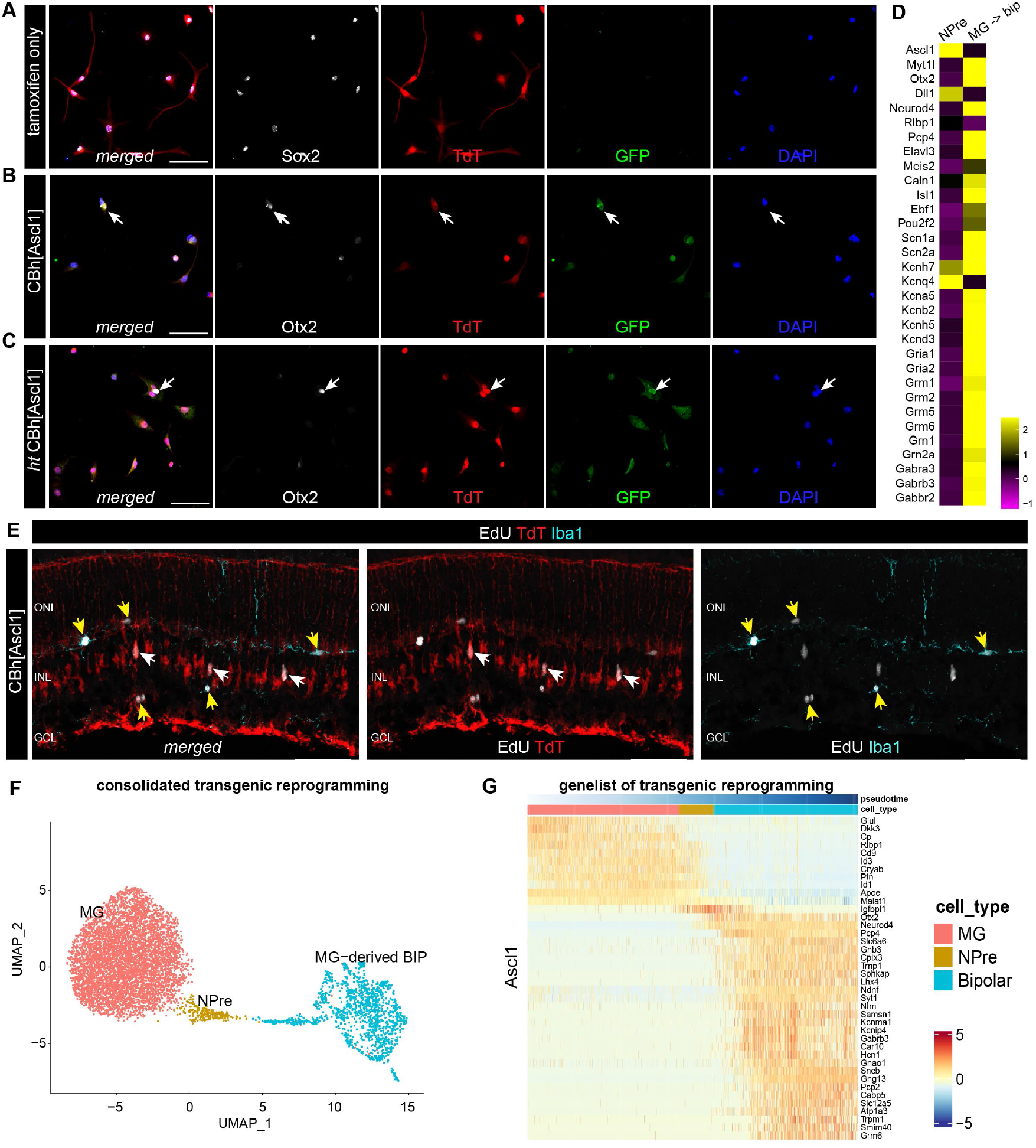
Histological and transcriptional profiling of AAV-mediated reprogramming. **(A-C)** fluorescence images of sorted lineage-traced cells on coverslips 24 hours post FACS split by condition showing merged and single channels of DAPI in blue, GFP vector reporter in green, TdT lineage tracer in red and glial marker Sox2 or bipolar marker Otx2 in white, **(D)** heatmap of average gene expression for selected neuronal genes in clusters NPre and MG-derived neurons following AAV-mediated reprogramming, **(E)** fluorescence images of retinal cross-section with proliferating lineage-traced MG (white arrows) and microglia (yellow arrows) with TdT in red, Iba1 in cyan and EdU in white, **(F)** UMAP of consolidated scRNA-seq data of Ascl1-mediated neurogenesis from transgenic animals (Glast-CreERt2 x LSL-tTA x tetO-Ascl1-GFP), **(G)** heatmap of top 40 differentially expressed genes across pseudotime trajectory from glial cell fate to neuronal cell fate based on scRNA-seq data **from (F); scalebar: 50 μm, ONL: outer nuclear layer, INL: inner nuclear layer, GCL: ganglion cell layer**.

**Fig. S3.**
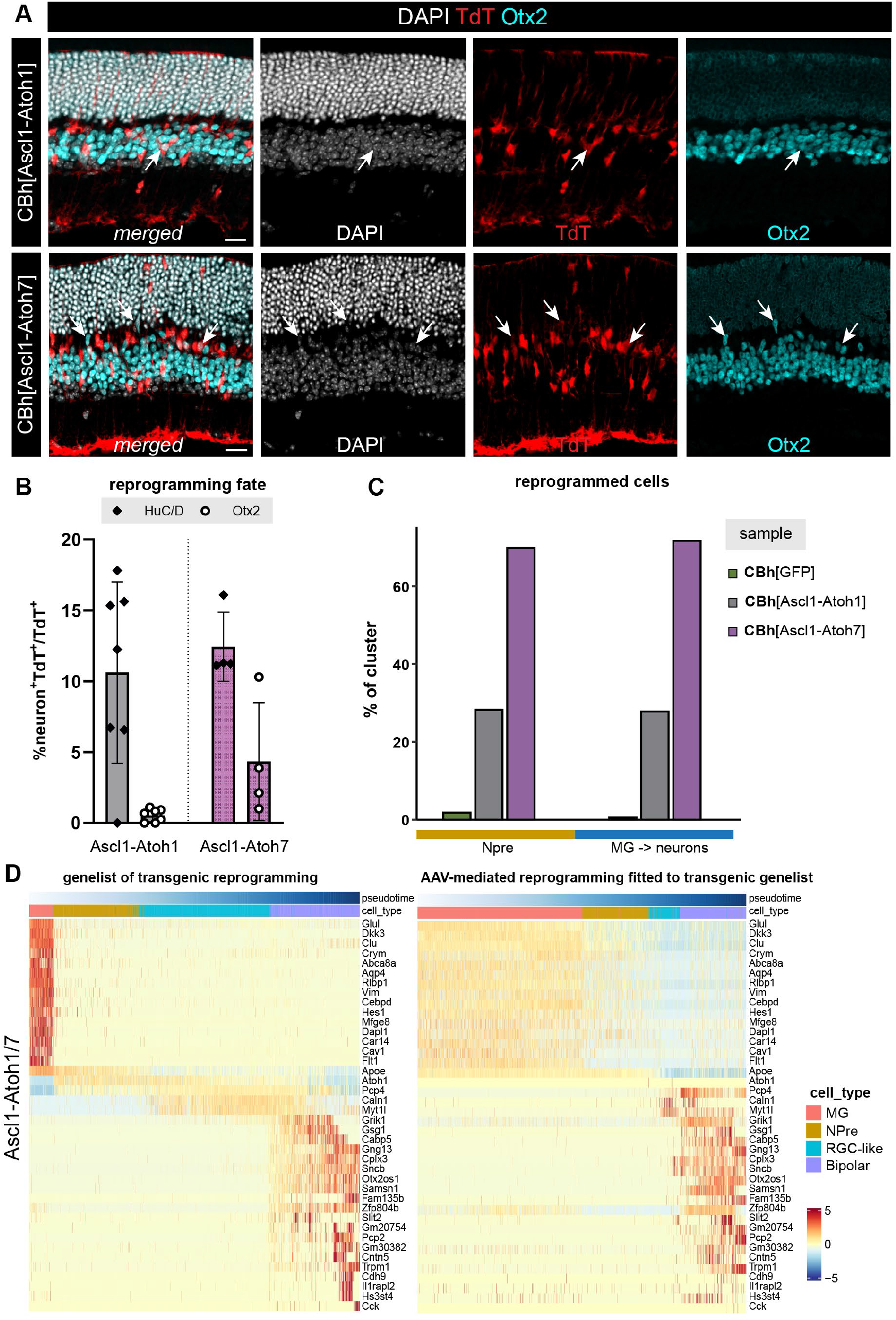
AAV-borne Ascl1-Atoh1/7 expression induces neurogenesis that phenocopies transgenics. **(A-A’)** fluorescence images of reprogrammed cells (white arrows) per condition showing merged and single channels of DAPI in white, TdT in red and Otx2 in cyan; scalebar for A-A’: 20 μm, **(B)** bar plot of MG reprogramming to distinct neuronal fates (HuC/D or Otx2) after AAV-mediated Ascl1-Atoh1 or Ascl1-Atoh7 expression counted as a percentage ratio of neuron^+^TdT^+^ over all TdT^+^ cells with each dot being a biological replicate (error bars: standard deviation), **(C)** barplot of the percentrage that each vector treatment contributed to the formation of reprogrammed clusters NPre and MG-derived neurons (MG→neurons) where each column represents a color-coded sample. **(D)** heatmap of top 40 differentially expressed genes across pseudotime trajectory from glial cell fate to neuronal cell fate based on scRNA-seq data from transgenic experiments, followed by heatmap of gene expression in scRNA-seq data from AAV experiments that follows the genelist generated from consolidated transgenic data for Ascl1-Atoh1/7 reprogramming.

